# Collective protection and transport in entangled biological and robotic active matter

**DOI:** 10.1101/2020.05.25.114736

**Authors:** Yasemin Ozkan-Aydin, Daniel I. Goldman, M. Saad Bhamla

## Abstract

Living systems at all scales aggregate in large numbers for a variety of functions including mating, predation, and survival. The majority of such systems consist of unconnected individuals that collectively flock, school or swarm. However some aggregations involve physically entangled individuals, which can confer emergent mechanofunctional material properties to the collective. Here we study in laboratory experiments and rationalize in theoretical and robotic models the dynamics of physically entangled and motile self-assemblies of centimeter long California blackworms (*L. Variegatus*). Thousands of individual worms form braids with their long, slender and flexible bodies to make a three-dimensional, soft and shape-shifting ‘blob’. The blob behaves as a living material capable of mitigating damage and assault from environmental stresses through dynamic shape transformations, including minimizing surface area for survival against desiccation and enabling transport (negative thermotaxis) from hazardous environments (like heat). We specifically focus on the locomotion of the blob to understand how an amorphous entangled ball of worms is able to break symmetry to move across a substrate. We hypothesize that the collective blob displays rudimentary differentiation of function across itself, which when combined with entanglement dynamics facilitates directed persistent blob locomotion. To test this, we develop robophysical blobs, which display emergent locomotion in the collective without sophisticated control or programming of any individual robot. The emergent dynamics of the living functional blob and robophysical model can inform the rational design of exciting new classes of adaptive mechanofunctional living materials and emergent swarm robotics.

**Significance Statement:** Living organisms form collectives across all scales, from bacteria to whales, enabling biological functions not accessible by individuals alone. In a few small cases, the individuals are physically connected to each other, forming to a new class of entangled active matter systems with emergent mechanofunctionalities of the collective. Here, we describe the dynamics of macroscopic aquatic worms that braid their long, soft bodies to form large entangled worm blobs. We discover that the worm blob behaves as a living material to undergo dynamic shape transformations to reduce evaporation or break-symmetry and locomote to safety against thermal stresses. We show that the persistent blob locomotion emerges as a consequence of physical entanglement and functional differentiation of individuals based on spatial location within a blob. We validate these principles in robophysical swarming blobs, that pave the way for new classes of mechanofunctional active matter systems and collective emergent robotics.

---

Active matter collectives consists of self-propelled individual units (living or artificial) that interact with each other to gain emergent functionality or to achieve common tasks (1–6). In these systems, the simple repeated interactions between the individuals and their environment can produce complex behaviors at the group level (3, 5). Depending on the type of interactions, the collective can display either fluid-like or solidlike properties (2). Fluid-like behavior is typically observed in unconnected individuals that avoid physical contact such as in flocking birds or schooling fish (5, 7–11). On the other hand, solid-like behavior is a consequence of physical contact between individuals such as in ants or bee self-assemblages (12–14). The latter type of entangled active matter aggregates enables the formation of large mechanically functional structures (bivouac, rafts, bridges etc.) that enable both new functionalities not accessible to the individual as well as enabling survival benefits to the collective, specially in harsh and adverse environmental conditions in which it is impossible for individuals to survive on their own (15–19).

In engineered systems, the emergent dynamics of active matter collectives has been explored in particles ranging in size from micrometers (active colloids) to centimeters (robots) (20–24). Specifically for collective swarm robotics, the majority of the past work has focused on mathematical modeling (25–29). These theoretical approaches often fail to adequately capture real-world physical interactions between individuals robots, which may critically influence the emergent collective behavior. Experimentally, although swarming systems have been successfully realized to collectively accomplish a common goal (30–32), each individual robot is equipped with costly and sophisticated sensors to leverage some degree of centralized control, which is subject to many limitations including low fault tolerance, scalability problems and design complexity (30). To overcome these limitations, researchers have proposed decentralized swarms, which eliminates the need for a central control unit, communication between individual agents and a priori knowledge about the environment (33). These decentralized swarm systems have only been demonstrated recently using physical entanglements, either magnetic (34) or geometric (35), that harnesses physical coupling between simple robots to yield task-oriented collectives capable of emergent functions.

In this study, we investigate worm blobs as an example of an entangled active matter where the long flexible bodies of blackworms (*L. Variegatus*) form transient links through braiding. The activity of individual worms in a blob enables it to self-organize and dynamically respond to changing environmental conditions. Depending on the type, history and gradient of the environmental stimulus (light, temperature, etc.), the blob can respond in a variety of ways. Here, we specifically focus on the evaporation and thermal response of worm blobs to understand why worms spontaneously aggregate into blobs and how they spontaneously move as whole. By developing simple robophysical blobs (36, 37) consisting of 3-link robots (smarticles) (35), we describe how variation of gaits and mobility of simple individuals in a physically entangled collective can lead to varying levels of locomotive performance, without the need for sophisticated central control systems.

## Results

### Worm blob as complex materials

Under threatening environmental conditions (evaporation, cold temperature etc.) individual worms spontaneously aggregate to form three-dimensional blobs that range from small collectives (N=10 worms) to large macroscopic entangled networks (N=50,000), both in water and in air (Fig.1,Movie S1). The blobs display non-Newtonian material properties (38), and can flow at long timescales (SI Appendix, Fig.S1D), while retaining shape to short time scale disturbances (SI Appendix, Fig.S1A,C). The underlying material timescale is set by the physical entanglement of the worm bodies through self-braiding (Fig.1). The blob formation and disintegration is reversible through ambient fluid temperature (SI Appendix, Fig.S1B). Increasing the fluid temperature ‘melts’ the blob (fluid-like phase), while decreasing temperature leads a more entangled (solid-like phase) (SI Appendix, Fig.S1B). We posit that the underlying principle for this emergent collective phase behavior lies in the changing activity of individual cold-blooded worms in response to external temperature and cooperative effects of active phase domains (38–41). At lower temperatures (*T <* 25^*o*^C), worms are less motile and elongated, while at higher temperatures (25 *< T* ≤ 30^*o*^C), they are more active and coiled up, with a peak activity at *T* = 30 ± 2^*o*^C (see SI Appendix, Fig.S2) (42). Thus, an entangled solid-like blob is observed at lower temperatures through slow-moving, elongated worms, while a disentangled fluid-like aggregate is observed at higher temperatures due to highly active, coiled-up worms, similar to motility-induced phase separations in active matter (41, 43).

**Fig. 1.**
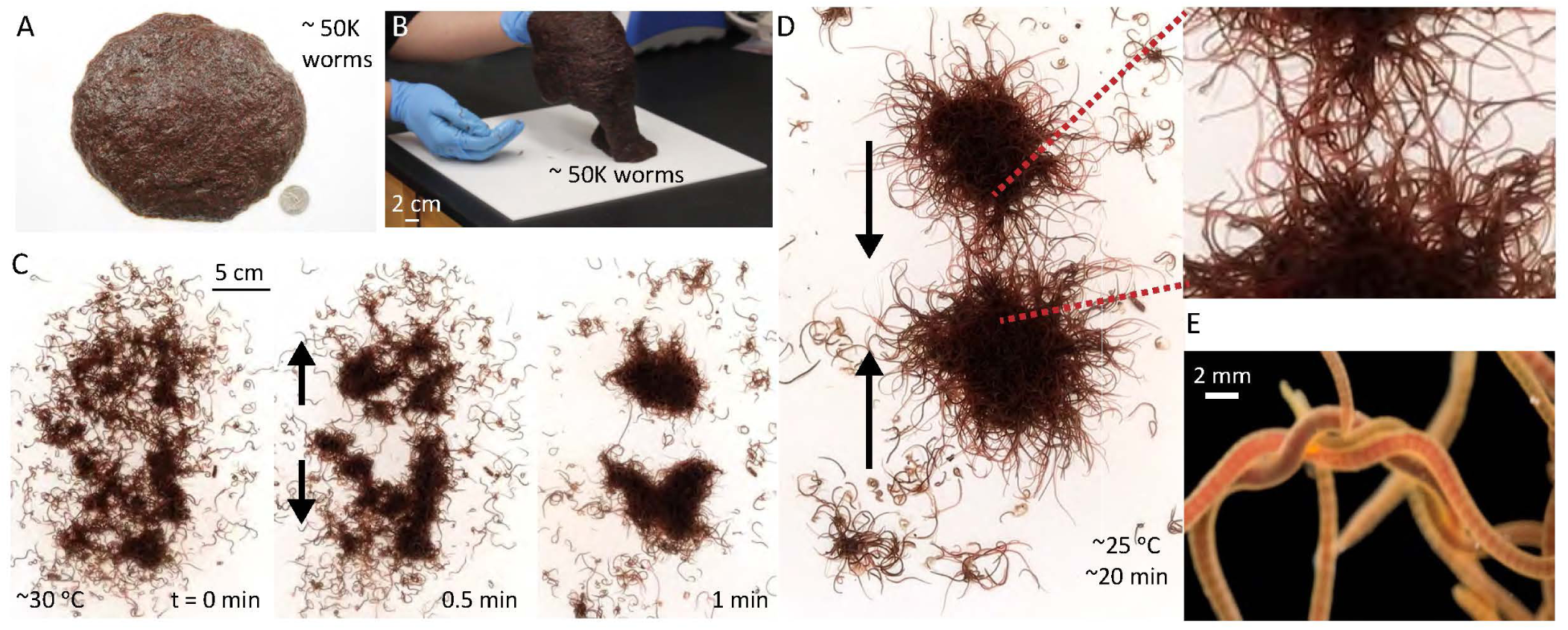
Worm blobs formed via physical entanglements. **A**. An entangled worm blob composed of ∼50,000 worms. **B**. The worm blob behaves as a non-Newtonian fluid, which can flow at long timescales and maintain shape as a solid at small timescales (Movie S1). **C-D**. Blob formation in water. The experiment starts at ∼30^*o*^C water in which the worms are mainly untangled with each other. As the water cools down to 25 ^*o*^C (using a Peltier cooler), the worms aggregate initially into two smaller blobs (t = 1 min), which ultimately merges to form one large blob (t = 20 min, see Movie S1). **E**. Close view of braid formation within a blob.

### Worms in a blob survive longer against desiccation

Worms are cold-blooded animals and their activities are greatly influenced by variations in temperature and light intensity of their surroundings (42, 44). Since blackworms naturally inhabit shallow aquatic regions (39), we hypothesize that forming a blob provides survival benefit to individual worms by reducing desiccation in air. To test this hypothesis, we expose blobs of different sizes (N=1-1000 worms, 10 replicates per condition) on a dry plate at controlled temperature and humidity (24^*o*^C, 48%), and track the projected blob area (A) using timelapse imaging over a few hours (see Movie S2). We observe that a single worm (*N* = 1) perishes in less than an hour, while worms in a blob (*N* = 1000) are alive even after 10 hours (see Movie S2). Additionally, the blobs are not static. Larger blobs (*N >* 20) undergo exploratory search modes for potentially favorable conditions, during which the blob area can almost double (40 mins, N=100) (see Fig.2A-B, Movie S2). This phase is followed by a shrinking mode during which the blobs become increasing spherical to minimize surface area to volume ratio. In contrast, smaller blobs (*N <* 20), monotonically shrink into spherical structures (Fig.2B). We note that this dynamic shape transformation behavior of worm blobs is reversible: addition of moisture once the blob has shrunk into a spherical shape, restarts the search mode (see Movie S2).

**Fig. 2.**
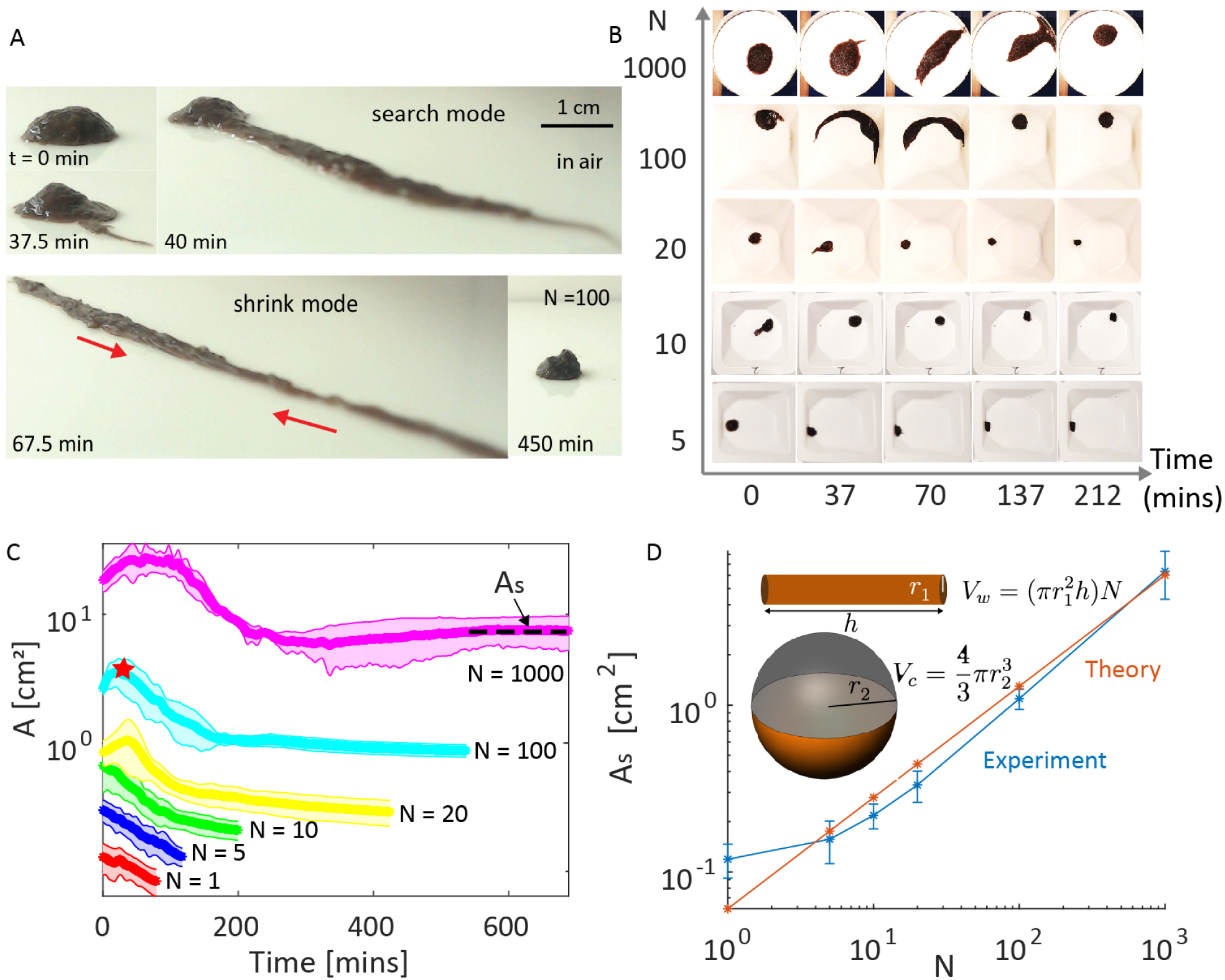
Evaporation response of the worm blob. **A**. When water is scarce, worms spontaneously form spherical blobs as a survival strategy to minimize evaporative losses. Time snapshot from the experiment (side view, N = 100 worms) for 450 minutes (Movie S2). The worms first undergo a stereotypical search mode for a water sources and after a critical time, spontaneously transition into a shrink mode to reduce surface area. **B**. The shape changes of the worm blobs in the air as a function of blob size (N=5, 10, 20, 100, 1000). See Movie S2 for the example experiments with N=1 and N=1000. **C**. Projected surface area (A) as a function of cluster size (N=1, 5, 10, 20 100 and 1000 worms, 10 replicates per condition) under controlled laboratory conditions (24^*o*^C, humidity 48%). Red star on the light blue curve indicates the time when the shrink mode starts. The worm blobs achieve a steady-state area (A_*s*_) indicated by a plateau in the curve, where the change in surface area is minimal (A_*s*_ = d(A)/dt*<* 1%). **D**. Comparison of experimental steady-state projected surface area (A_*s*_) (blue) with theoretical estimation of surface area (red) across three orders of magnitude of blob size (N) reveals good agreement between model and experiments.

To support the hypothesis that the worm blobs are reducing evaporative losses through minimizing surface area, we quantify the steady-state projected surface area (A_*s*_, defined the surface area is invariant, i.e. d(A)*/*dt*<* 1%) as a function of blob size *N* as shown in (Fig.2C-D). Theoretically, we estimate the volume of *N* worms as 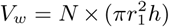, where each worm is assumed to have cylindrical body of radius of *r*_1_ and length of *h*. If these *N* worms minimized surface area to form a sphere, then the theoretical projected surface area of the hemisphere is given by 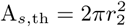, where 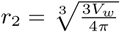 is the equivalent radius. Indeed, the experimental data and simple theoretical model are in good agreement over three orders of blob size *N*, validating the hypothesis that worms blobs formation spherical shapes to reduce evaporation losses (Fig.2C-D). Beyond *L. Variegatus*, other annelids that we tested, such as *L. terrestris* and *E. fetida*, (SI Appendix, Fig.S3, Movie S2) as well as past observations on nematodes (*C. elegans*) (45–49), suggest forming entangled collectives may be a general biological strategy to survive desiccation for extended periods by these organisms in fluctuating arid environments.

### Emergent locomotion of worm blobs

Since the worms are sensitive to temperature (as described earlier) as well as light (39, 40), we next investigate the emergent behavior of the worm blobs to a combination of light and temperature cues in a custom setup as shown in Fig.3A (see details in SI Appendix, Fig.S4). In laboratory cultures at constant temperature (∼15^*o*^C), we observe that the worms form tightly entangled blobs at high light intensities and form loose dispersions in the dark. A sudden increase in light intensity results in a rapid blob contraction (33±6%) reduction, after 15 hours in dark) in a short duration (*t <* 5 s, SI Appendix, Fig.S5, Movie S3), which is expected as individual worms are known to exhibit a rapid contractile escape response to shadows and photic stimulations arising from movements of overhead predators in the water (50). Thus, bright light serves as a cue for worms to aggregate and entangle tightly.

**Fig. 3.**
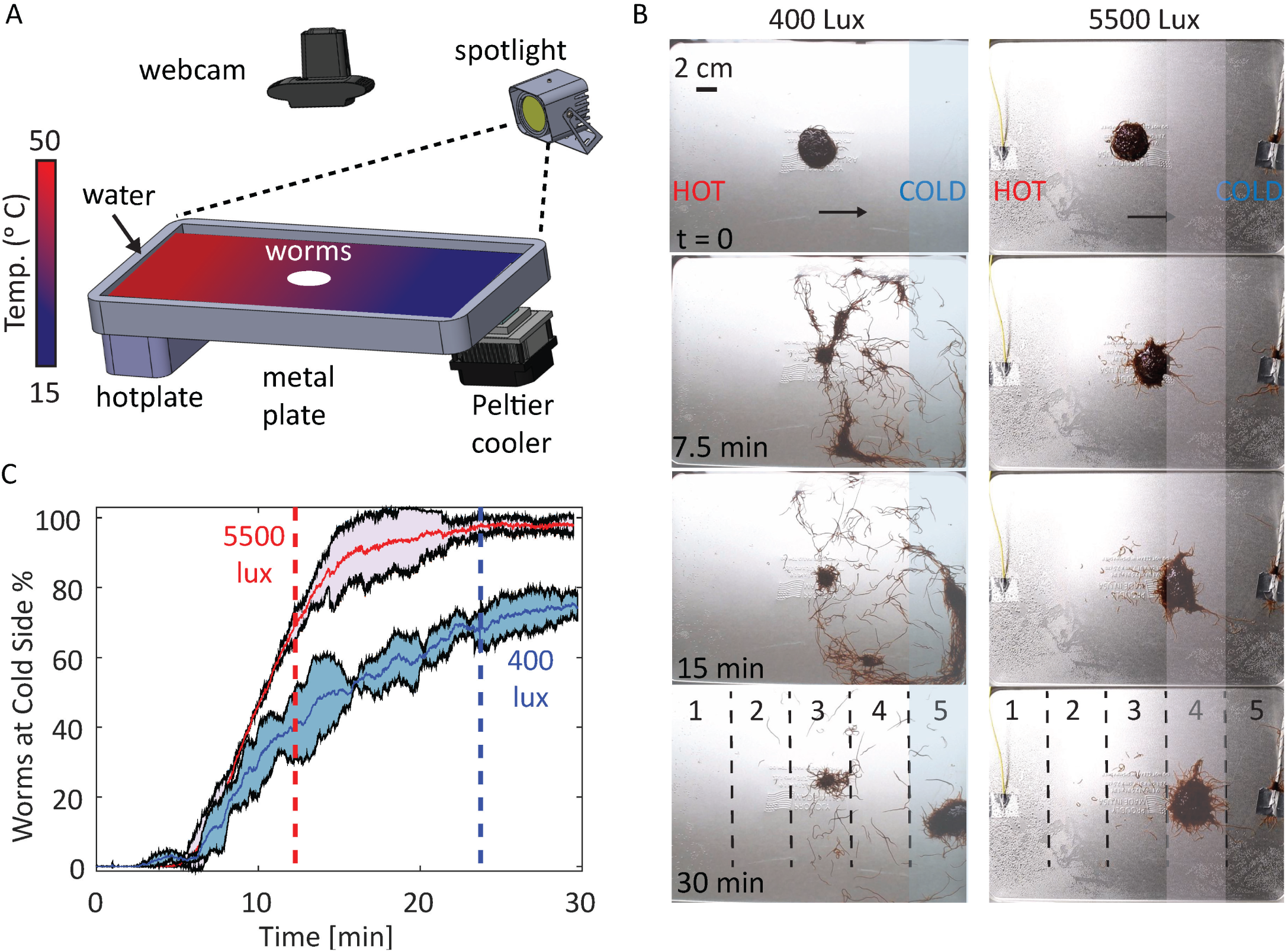
Worm blobs exhibit emergent locomotion under thermal gradients. **A**. Schematic of custom built experimental setup to study worm blob locomotion under thermal gradients and different light conditions. The worm blob is placed (*N* = 600 worms.) into the center of a metal plate (30×20×5cm) filled with water. We establish thermal gradients on the surface of the plate by setting the temperature of the cold and hot side to 15^*o*^ and 50^*o*^C, respectively (see SI Appendix, Fig.S4 for the details of the setup). Color bar represents the temperature of the water. **B**. Time snapshots (t=0, 7.5, 15 and 30 min) from the thermotaxis navigation experiments under room light (400 lux, left) and spotlight (5500 lux, right). In both cases, the worms exhibit negative thermotaxis, but under low light conditions, they move individually, while under high light intensities, they move collectively as a blob. Dashed lines divide the plate into five equal areas for tracking movement of worms across the plate. For both experiments, overlap space time heat maps of worm locomotion are shown in SI Appendix, Fig.S6,7. **C**. During the same duration, by moving together as a blob, *>* 90% of the worms make it to the colder side (zone 4), while moving individually *>* 70% of the worms make it to the cold side (zone 5). Dashed lines (red, 5500 lux and blue, 400 lux) show the time when same amount of the worms (70%) reach the cold sides in both experiments.

Under room light intensity (∼400 lux) if we next expose the blob to an approximately linear temperature gradient (see the experimental setup in Fig.3A), the blob dissipates and worms individually crawl to the cold side (Fig.3B left column, Movie S3,SI Appendix, Fig.S6). Without changing the temperature gradient when we increase the light brightness to 5500 lux, we discover a surprising behavior: worms exhibit emergent locomotion to collectively move as a blob towards the cold side (negative thermotaxis, Fig.3B right column, Movie S3, SI Appendix, Fig.S7). In both cases, majority of the worms (>70% for 400 lux and >90% for 5500 lux) are successfully able to move to the towards the colder side, with a similar average speed (0.6 ± 0.1 cm/min, calculated when 70% of worms reach cold side) as shown in Fig.3C.

To examine if moving together as a blob vs. moving individually conferred additional benefits, we utilize a quasi-2D experimental apparatus, which facilitates easier worm tracking (see SI Appendix, Fig.S8A-B). We find that worms crawling individually can move at faster speeds to the cold side (N=10, *v*_*s*_ =1.2*±*0.3 cm/min) as compared to worms in a blob (N=80, *v*_*b*_ =0.4*±*0.1 cm/min). However, moving as a blob despite being slow, enabled all the worms to be transported safely to the cold side. Not all worms moved to safety when they crawled individually (SI Appendix, Fig.S8C). Thus for an individual worm, being in a blob confers multiple survival benefits: reduced evaporation when water is scarce and reliable transport to safety when the environmental temperature becomes fatal.

### Mechanism of emergent blob locomotion

How does an entangled worm blob spontaneously break-symmetry and move? We hypothesize that the locomotion of the blob towards the cold side emerges as a consequence of the individual worm response to temperature. To test this hypothesis, we recorded close-up videos of small worm blobs (*N* = 20) while moving under a thermal gradient (Fig.4A). Using their circular and longitudinal muscles, the worms can apply contractile pulling forces along the length of their body to crawl forward (51). As described earlier, the the motility (and activity) of the worms increase at higher temperatures SI Appendix, Fig.S2, (42). Thus, depending on the position in a blob (front or rear), individual worms encounter different thermal stimuli. We observe that worms facing the cold side (leading edge) act as pullers and use their elongated bodies to apply slow, periodic pulling forces on the blob (Movie S4, Fig.4B-C). In contrast, the worms closer to the hot side (trailing edge) are more coiled up and exert minimal traction forces in the direction of locomotion. However, the coiling behavior of the trailing edge worms acts to lift up the back of the blob, potentially to reduce friction (Fig.4C).

**Fig. 4.**
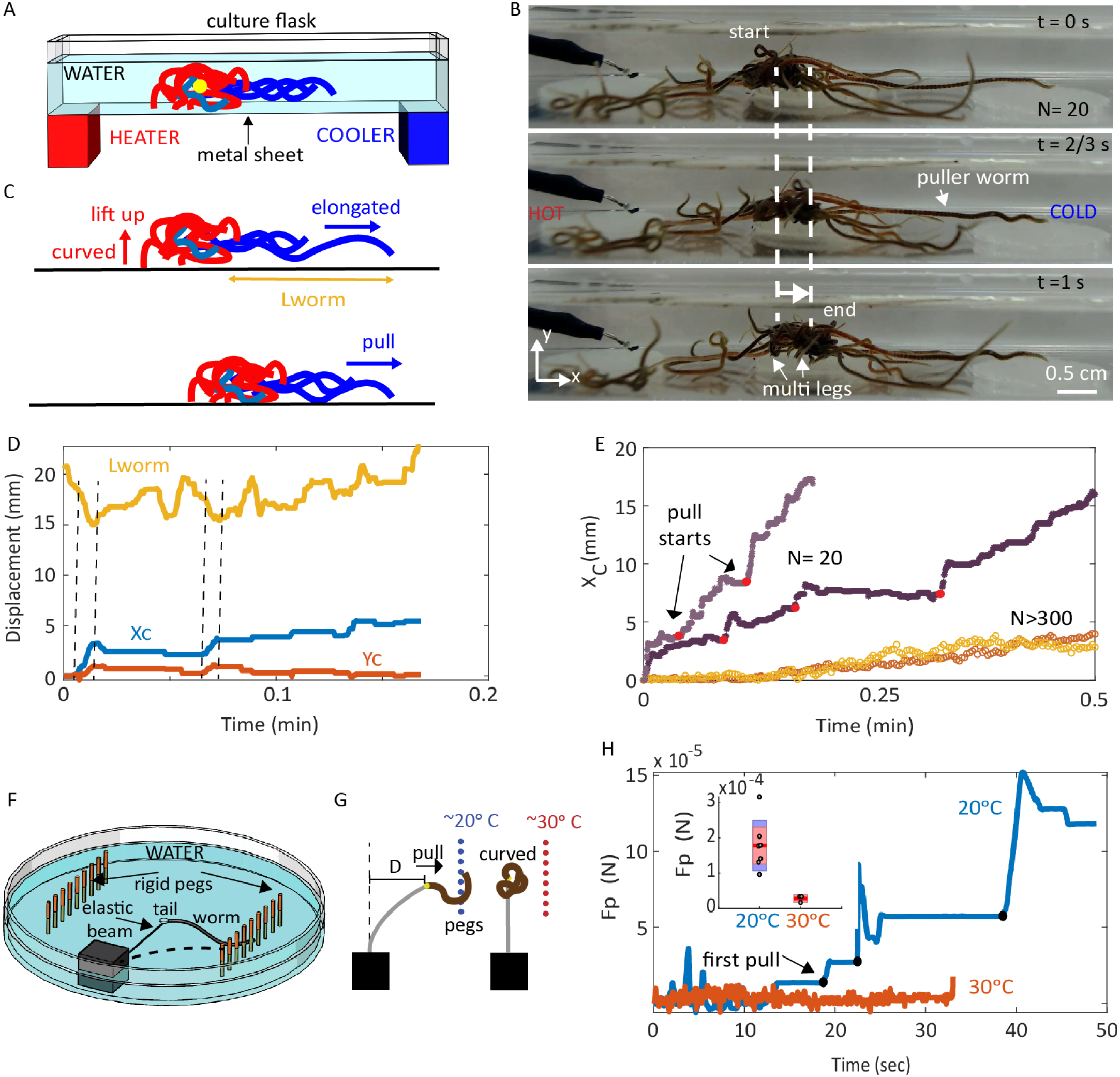
Mechanism of emergent locomotion of a worm blob through differentiation of activity. **A**. Schematic of the experimental set-up with temperature gradient. The yellow dot shows the center of blob (CoB). **B**. Snapshots from the quasi-2D experiment, where a small worm blob (N=20) exhibits negative thermotaxis, crawling towards the cold side (right) in response to a temperature gradient (Movie S4). **C**. Proposed mechanism of how an entangled worm blob breaks symmetry to exhibit directed motion. The worms blob moves through differentiation of activity: worms at the leading edge (blue) act as pullers, while worms at the trailing edge (red) curl up and lift the blob to potentially reduce friction. **D**. Changes of the length of the leading edge puller worms (*l*_worm_) and the center of blob position, X_*c*_ (blue) and Y_*c*_ (red), as a function of time. The maximum horizontal displacement coincides with pulling (shortening of worm length) and lifting events as indicated with dashed vertical lines. **E**. Horizontal displacement of center of blob for N=20 (dark and light purple) and N*>*300 (dark and light yellow) in response to thermal gradients. Red dots indicates where the pulling events start (Movie S4). **F**. Schematic of the experimental setup to measure pulling force of individual worms. Series of rigid pegs are mounted on a plastic petridish (100 × 15 mm) and the tail of the worm is glued to force calibrated elastic beam SI Appendix, Fig.S9. By measuring the deflection of the beam by the worms, pulling force is estimated. **G**. Illustration describing the observed behavior of worms during measurements at 20^*o*^C (blue) and 30^*o*^C (red) as shown in Movie S4. **H**. Force measurements for single worms in cold (20^*o*^C, blue) and hot water(30^*o*^C, red). The black dots on the blue curve indicates the start time of successive pulling events by worms as seen in Movie S4. Inset shows the mean and standard deviation of the maximum pulling forces in cold (5 trials) and hot water (3 trials).

We quantify both the change in the length of the leading worms *l*_worm_ and the displacement of the center of blob (X_*c*_, Y_*c*_) in Fig.4D. The large forward displacements occur in sync with both, contraction of the puller worms (0.15±0.06 *l*_worm_) and vertical lifting of the blob (by trailing edge worms) as indicated by the vertical dashed lines in Fig.4C). The pulling force mechanism is evident by distinct pull events during locomotion of small blobs (*N* = 20) across a substrate (Fig.4E). This proposed mechanism of functional differentiation of worms in a blob into puller worms at the leading edge and friction reducing worms indeed also holds for larger blobs (N*>*300), as observed in the supplementary video (Movie S4). We note that the displacement (X_*c*_) for larger blobs is smoother compared to the intermittent pulling events of smaller blobs Fig.4E, potentially due to larger number of puller worms working in tandem (Movie S4).

To test if a single worm can generate the necessary force to pull a blob, we tether individual worms to a calibrated cantilever beam and measure the pulling force exerted by a single worm on a rigid peg (Fig.4F, SI Appendix, Fig.S9). At low temperatures (20^*o*^C), the worms are more elongated and use their bodies to exert large pulling forces (*F*_*c*_ = 178.2±52.5 *µ*N, ∼2.5 times weight of a single worm). While at high temperatures (30^*o*^C) the forces were nominal *F*_*h*_ = 28.0 ± 10.9 *µ*N (see Fig.4G-H). Assuming a low coefficient of static friction for wet acrylic (*µ*_*s*_ = 0.3, substrate used in our experiments) and a blob mass of *m* = 0.14g (N=20 worms), we estimate 2-3 individual worms could generate sufficient traction force (*F* = *µ*_*s*_mg ∼412 *µ*N) to move the small blob. This estimate qualitatively agrees with experimental observations, where individual worms seem capable of moving the blob (Fig.4C, Movie S4). Thus, the locomotion in the blob emerges through differential activities of the individuals in the front and rear, in response to temperature.

### Robophysical blobs

Here we develop a robophysical blob to validate our hypothesis that an entangled collective can exhibit emergent locomotion through two basic principles: mechanical interactions (entanglements) and differentiation of roles in the collective. The robotic blobs consists of six 3-link, 2 revolute joints, planar, smart, active particles (smarticles (35)) equipped with two light sensors as shown in Fig.5A. Unlike a previous study of the collective smarticle locomotion (35) where weak entanglement was achieved by shape changes of the arms and an external ring confinement, here we purposefully achieve strong entanglements through meshes and L-shape pins on the robot arms (Fig.5B). The length (width = 1 cm, leg length = 0.5 cm), shape and orientation of the pins were chosen so that the arms easily attach and detach to the mesh and form an entangled robotic blob mimicking the worm blobs (see Fig.5C).

**Fig. 5.**
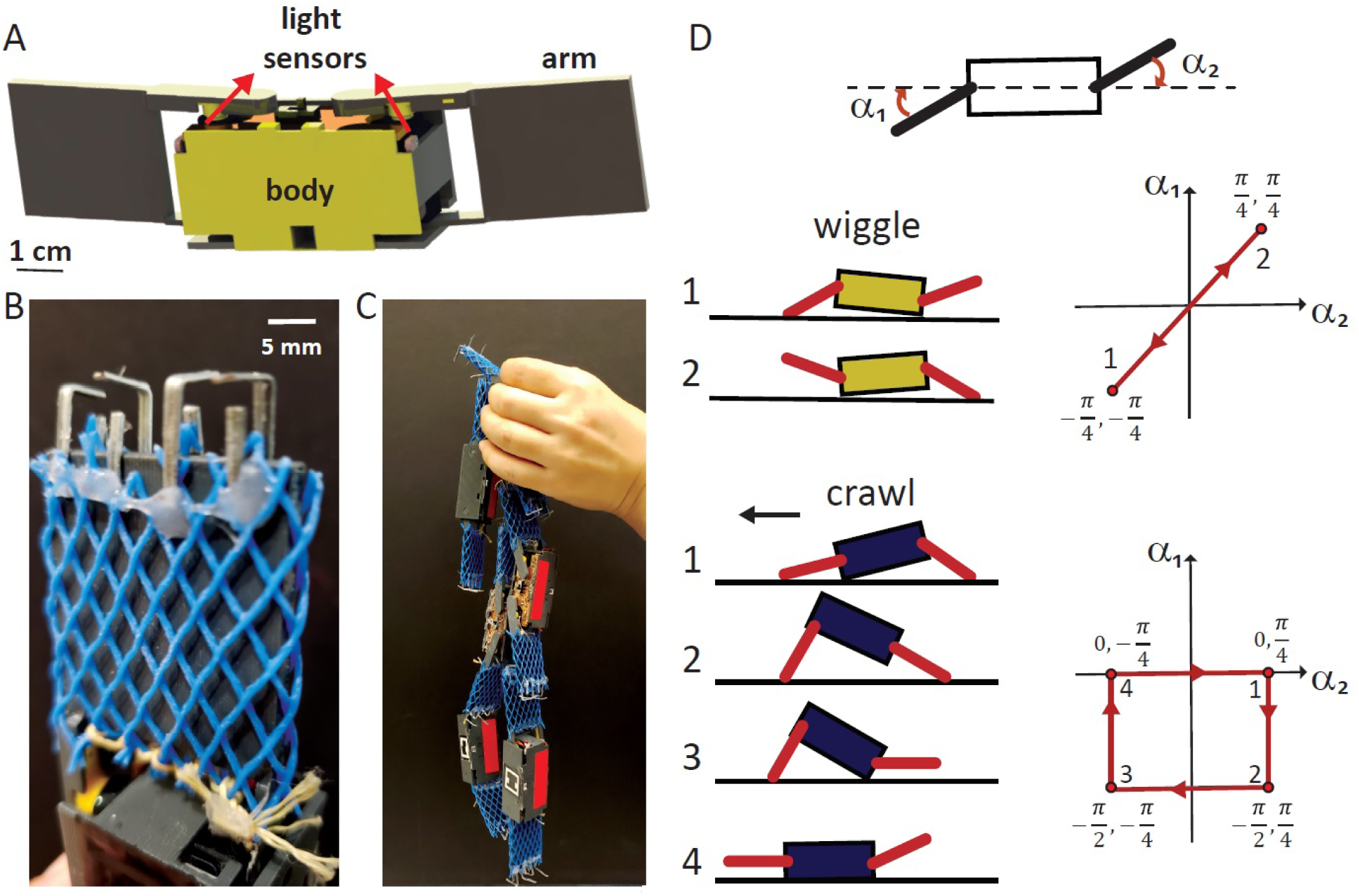
Robophysical model of a worm blob consisting of small 3D printed robots. **A**. Each robot is a 3-link, planar robot equipped with two photoresistors and the arms are connected to the body via servos controlled by Arduino Pro Mini (35). **B**. To enhance the physical entanglement of the robots, the arms are covered with a plastic mesh and L-shape pins are inserted to the edge of the arms. **C**. Six robots entangle to each other to form a bioinspired robotic blob. **D**. Motion sequences of two gaits as a function of arm angles (*α*_1_ and *α*_2_), wiggle (top) and crawl (bottom). We define *α*_1_ *>* 0 when it is above the centerline and *α*_2_ *>* 0 when it is below the centerline. The arrows show the direction of the arm movement sequence. Note that, the wiggle gait does not lead to forward motion while the crawl does (to the left).

To achieve autonomous collective locomotion in response to a light stimulus, we program each robot with two behaviors: ‘wiggle’ and ‘crawl’ as a function of their arm angles *α*_1_ and *α*_2_ (Fig.5D, Movie S5). In the wiggle gait, {*α*_1_, *α*_2_} = [(−*π/*4, −*π/*4), (*π/*4, *π/*4)] such that the robot arms swing up and down (out-of-phase with each other) 45^*o*^ from the center line of the body Fig.5D. While in the crawl gait, *α*_1_ amd *α*_2_ extend up to 90^*o*^, following a sequence as shown in Fig.5D. We note that only the crawl gait leads to forward motion towards the light source (positive phototaxis). The robots switch between these two gaits by sensing the light intensity of the environment using two optical sensors on their body. The wiggle gait is activated when the light intensity is below a threshold (*<*200 lux, dark), while crawl gait is activated above a certain threshold (*>*800, light). Thus similar to the biological system (worms) that have have different activity levels at low and high temperatures (thermotaxis), the robotic models have two different gaits in response to light intensity levels (phototaxis).

### Emergent locomotion of robotic blobs via gait differentiation

We confine six robots in a 2D arena (l=60 cm, w=5 cm), where the robots touch the sidewalls and half of the bottom surface is covered with a mesh to increase substrate friction (Fig.6A). A light source is mounted on one side of the box to activate the gaits as described earlier. A stereotypical configuration of the robotic blob is shown in Fig.6A, where the leading robot (blue) executes a crawl gait towards the light source and acts as puller for the entire collective. The bottom robots (yellow) robots can potentially execute a wiggle gait to reduce the contact area of the blob.

**Fig. 6.**
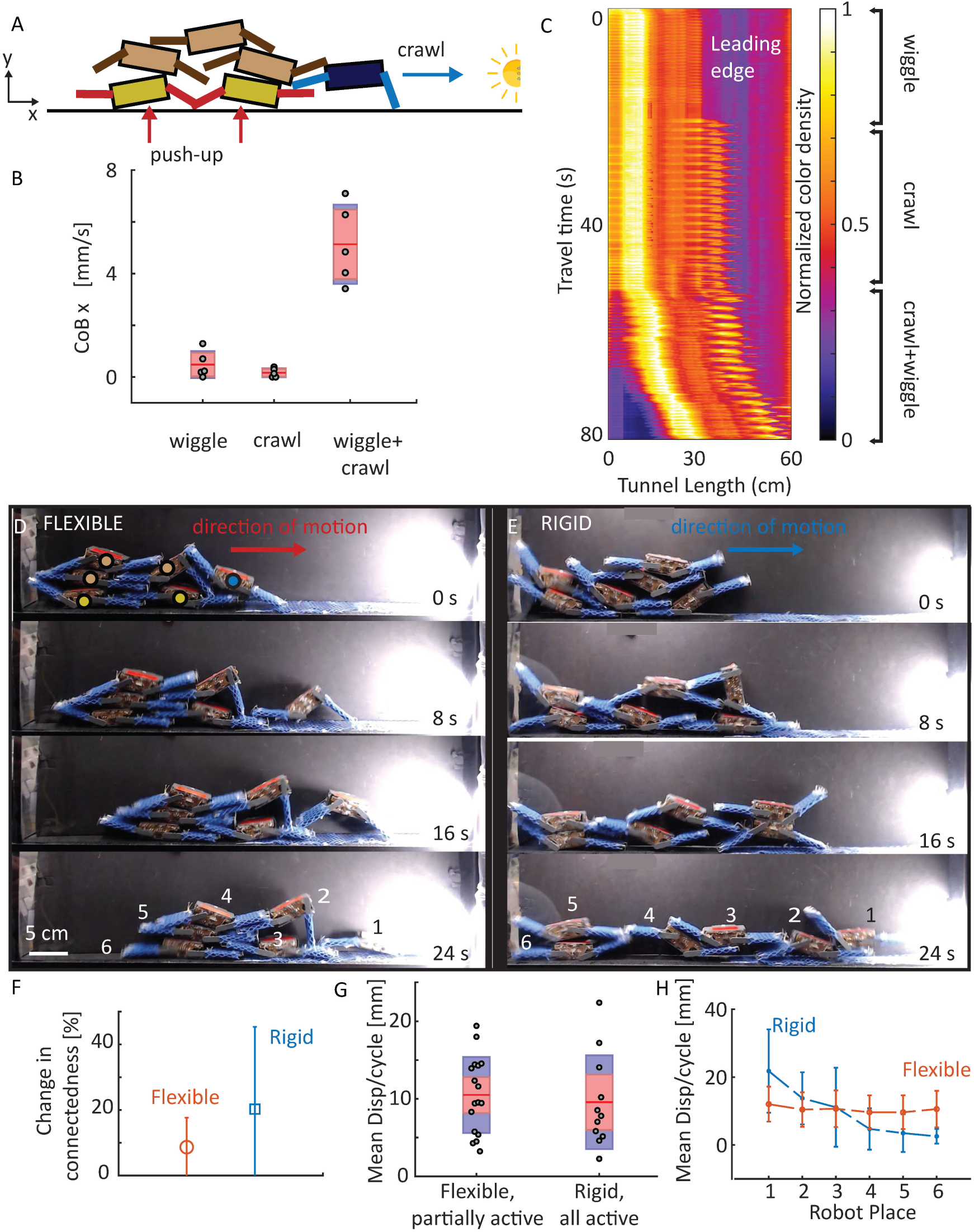
Mechanical interactions and collective movement of a physically-entangled robotic blob. **A**. Schematic of the robotic test arena, where the first half of the surface is covered with a mesh to increase substrate friction and a light source is used to activate different gaits (phototaxis). The leading robot (blue) crawls towards to light, bottom robots (yellow) wiggle to push up the center of mass of the blob. Other robots (brown) remain completely or partially immobile during the experiment by keeping arms either rigid or flexible. **B**. The comparison of the mean and standard deviation of a displacement of the CoB of the robotic blob over 6 test runs for 3 conditions: only bottom yellow robots wiggle, only leading blue robot crawls, and both blue robot crawls + bottom yellow robot wiggle simultaneously (Movie S5). The red lines represent the mean value, the red area represents the 95% confidence interval, and the blue area represents the mean *±* one standard deviation. **C**. Combined space-time plot of the 3-condition experiment shown in Movie S5, where only the bottom yellow robots wiggle (0-20s), only leading blue robot crawls (20-50s), and both yellow robots wiggle + leading blue robot crawls at the same time (50-80s). We note that only when differentiation of gait occurs in the entangled robotic blob (crawl + wiggle), do we observe directed motion towards the light. Color bar shows the normalized color density of the robots and the setup. **D**. Snapshots from the experiment where the robots with brown dots are inactive and their arms are kept flexible. Other robots changed their gaits according to light intensity thresholds described in the main text (Movie S5). **E**. Snapshots from the experiment where all the robots are active. At intermediate light intensity (200 *−* 800 lux), the robots become inactive and kept their arms rigid (Movie S5). **F**. Change in the number of robots in contact with each other, at the beginning and at the end of the experiments. For the flexible case shown in D (n=17 trials, red) and rigid case shown in E (n=11 trials, blue). **G**. Mean displacement of all the robots per cycle in a run for the flexible (n=17 trials) and rigid (n=11) cases. **H**. Mean displacement of the individual robots in a run for the flexible and rigid cases.

First, we demonstrate that the differentiation of roles in a robotic blob is critical for collective transport. By sequentially switching the gaits of the individual robots (if only the bottom robots wiggle or if only the pulling robots crawl) the blob does not move (Fig.6B, Movie S5). For persistent directed collective motion, both the puller (crawl) and friction reducing robots (wiggle) needs to be simultaneously run, but need not have their gaits synchronized in any way (Fig.6C, Movie S5). Thus, by simply programming gait differentiation across an entangled robot collective, we further support the mechanism proposed for the emergent locomotion of the biological worm blob.

Next, we conduct an additional set of experiments to investigate the parameter space that affects the collective transport dynamics. By altering the states of the top robots (brown shown in Fig.6A), we describe how the swarming behavior of the robotic blob can transition from moving collectively as a group to individually. In the first case referred to as ‘flexible’, the top (brown) robots are inactive, and thus their arm positions change only through their physical interactions with neighboring robots. The remaining robots are active and change their gaits according to the light intensity detected by the sensors as described earlier. Thus, robots at the bottom execute a wiggle gait since they are covered by other robots and shadowed from the light, while the leader robots execute a crawl gait due to exposure to the light source. Effectively, this reproduces the same result as the wiggle and crawl mode described above in Fig.5C, albeit dynamically as the robots spontaneously switch their gaits based on their spatial location in the group, resulting in the robots moving together as a group towards the light Fig.5D, Movie S5).

In the second case referred to as ‘rigid’, initially all the robots are active, but we define an additional state where robots with intermediate light intensity (200 − 800 lux) become inactive and keep their arms rigid (inflexible). Now the robots slowly disentangle and crawl individually towards the light (Fig.6D). Thus, we observe that entanglement is key for collective transport as in the flexible case, there is very little change in connectivity between the robots (8.7 ± 9.0 %), while in the rigid case the robots go from highly connected to spread out (20.1 ± 25.1 %) as shown in Fig.6F. We note that the mean displacement of the entire group is quite similar in both cases (flexible: 10.5 ± 4.9 mm, rigid: 9.6 ± 6.0 mm), however the mean displacement differs significantly for individual robots in the rigid case as only 1-2 of robots successfully make it all the way to the light source (Fig.6F). Thus, similar to the biological systems where the worms can move either individually or as a blob with varying levels of functional benefits, we demonstrate that the robophysical model of a worm blob can be tuned through simple gait strategies and dynamic local interactions to achieve different levels of collective locomotion performance.

## Conclusion

We have performed the first functional study on a biological system consisting of a multitude of worms,i.e. worm blob, to investigate the fundamental mechanisms behind the emergent physical adaptability, mechanofunctionality, and locomotion of the entangled collective. We studied systematically how the worm blob reacts to different environmental stresses depending on the type, history and intensity of the associated perturbations. We found that a worm blob can collectively move through the use of physical entanglement and differentiation, which allows an emergent behavior even in the absence of centralized control. To validate our hypotheses on the collective locomotion of the worm blob, we used a robotic blob made of six small, 3-link robots that can entangle to each other with the help of mesh covered pinned arms. Our robophysical system enabled us to investigate other parameters such as the gaits, activity and flexibility of the individuals that are challenging to directly test in the living system.

For robotics, the creation of a coherent swarm of simple robots has been a dream of roboticists for years, and our robotic system is part of an emerging trend in leveraging mechanics and physics to perform collective tasks in a decentralized way (34, 35) rather than the traditional algorithm-based and centrally-controlled approach to swarms (30, 52–58). For biology, the worm blobs hold exciting potential to inspire adaptive active materials as well as advance our understanding of emergent biomechanics of living collectives (15–19). We note that, to the best of our knowledge, the only other entangled assemblage capable of emergent motility occurs at cellular scales, where the amoeboid cells of the slime mold *D. discoideum* form a motile slug synchronized by cAMP waves (59). At larger length scales, almost exclusively all known examples of functional self-assembled structures (bivouacs, rafts, bridges etc.) are observed in insect societies (Phylum: *Arthopods*), which although can adapt and reconfigure, do not exhibit emergent locomotion of the whole entangled collective (12). Thus this report contributes the first discovery of a physically entangled and self-motile self-assemblage in a non-Arthopod multicellular organism. While beyond the scope of this paper, we have so far observed an impressive array of behaviors of this exciting system, including food foraging by the blob using the individual worms as appendages and locomotion in unstructured three-dimensional environments, opening up further discoveries of emergent behaviors in a seemingly mundane blob of squishy worms.

## Materials and Methods

### Animal Experiments

*Lumbriculus variegatus* was obtained from Aquatic Foods & Blackworm Co (CA, USA). The worms (length=2.5 ± 1 cm, diameter=0.7 ± 0.25 mm, mass=7.5 ± 3 mg) were cultivated in a box (35×20×12 cm, 25 gr worm per box) filled with distilled water (h = ∼2 cm) at ∼15^*o*^C for at least three weeks prior to experiments. We feed the worms with tropical fish flakes once a week and change the water one day after feeding them. Studies with *Lumbriculus variegatus* do not require approval by the institutional animal care committee.

### Analysis of the data

All the animal data were analyzed using the MATLAB Image Processing Toolbox.

## ACKNOWLEDGMENTS

We thank Will Savoie, Ross Warkentin, Max Seidel and Meredith Caveney for early design of smarticles. This work was supported by Soft Matter Incubator (SMI) Seed Grant Program of Georgia Tech. M.S.B acknowledges support from NSF (no. 1817334 and CAREER grant no. 1941933).

