## Supplementary Information for "Collective protection and transport in entangled biological and robotic active matter"

May 24, 2020

#### 1 List of supplementary videos

1. **Movie 1: Entanglement, blob formation and material properties of the worm blob** (1) Blob formation in water starting with untangled state ($30^oC$). (2) $\sim$50K worms form a viscoelastic blob. (3) Viscosity, elasticity and reversible state transition of the worm blob.
2. **Movie 2: Evaporation response of a worm blob.** (1) A single and thousand worms respond the desiccation. (2) Search and shrink behavior of the worm blob ($\sim$100 worms) under room light. (3) Shape changes of the worm blob in water and in air under spotlight. (4) Reversible evaporation response of the worm blobs under room light. (5) Entangled aggregation behavior of other annelids (*L. terrestris* and *E. fetida*)
3. **Movie 3: Phototaxis and thermotaxis responses of a worm blob.** (1) Temperature response of a single worm. (2) Contraction of a worm blob under different light history. (3) Collective locomotion under thermal and light stress.
4. **Movie 4: Locomotion mechanism under thermal stress via differentiation in function of regions of a blob and entanglement.** (1) Pull-push mechanism of a worm blob ($\sim$ 20 worms) under thermal stress. (2) Locomotion of a worm blob ($\sim$300 worms) under thermal stress. (3) Pulling forces of a single worm in cold and hot water.
5. **Movie 5: Mechanical interactions and collective movement of a physically-entangled robotic blob.** (1) Crawl and wiggle gaits. (2) Differentiation mechanism. (3) Moving as a blob or individually depending on the activity of the robots.

### 2 Supplementary figures

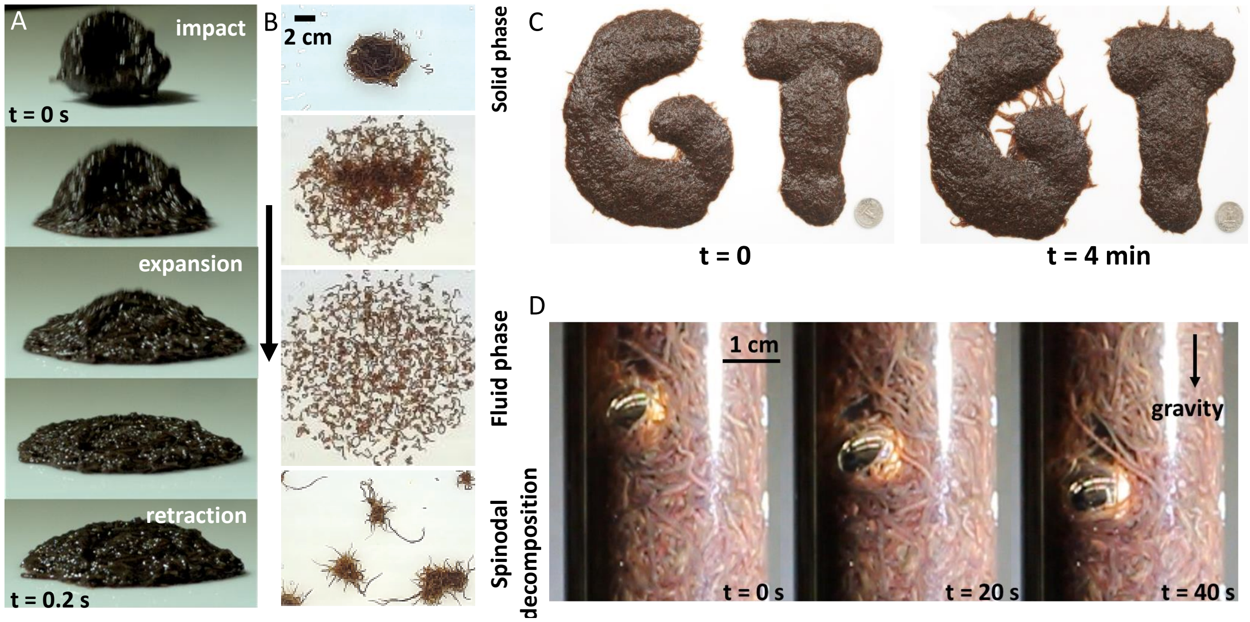


Figure 1: **Material properties of the worm blob– A.** The worm blob (consists of 5gr. alive worms) was dropped from 30 cm height. When the droplet comes into contact with a hard surface, it expends and and retract in 200 ms. The resulting worm aggregation exhibits elastic behavior. **B.** Solid and fluid-like behavior of worm blob in water. Immediately, after we put the worm blob into the water (temperature is about $\sim 32^{o}$C), the individual worms in the blob disentangles and changes the state of the blob from solid to fluid. The state transition is reversible. After adding cold water, they started to aggregate and create small blobs in 15 min (bottom). **C.** 3D living sculptures with 50k worms. **D.** The steel sphere of diameter 1.2 cm with a density of $\rho_{sphere} \approx$7.5 g cm$^{-}$3 is placed on top of a worm aggregation ($\rho_{worm} \approx$ 1 g cm$^{-}$3) inside a glass measuring tube (d = 3.5 cm). The gravitational acceleration is g = $10^3$cm s$^{-2}$. The sphere falls through the aggregation along vertical direction with a speed about v$\approx$0.025 cm s$^{-1}$. This experiment shows the viscosity of the worm blob (see Movie 1).

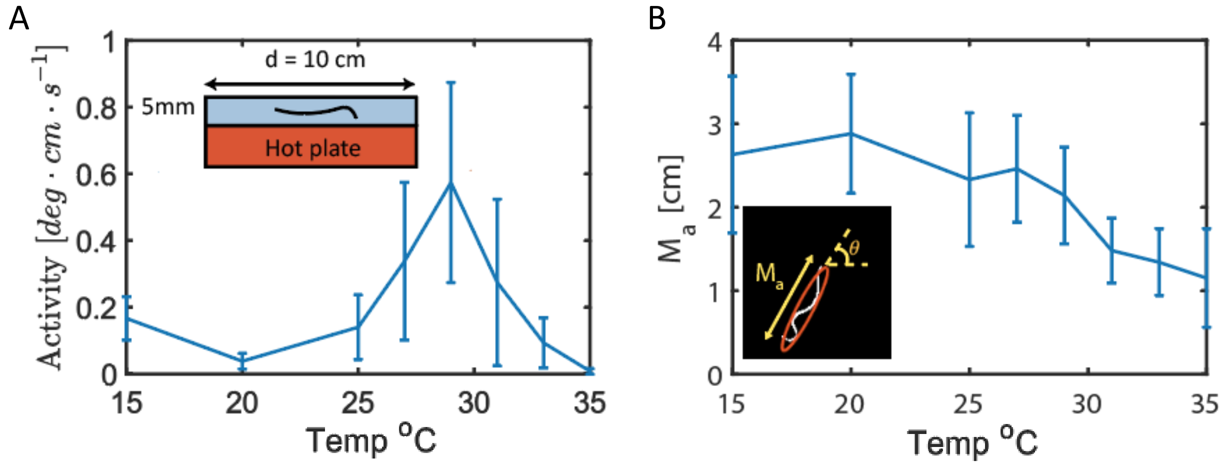


Figure 2: **The temperature response of a single worm**– Single worm was placed into petridish (d = 10 cm) filled with water (h = 5mm). The temperature of the hot plate was adjusted and we waited 5 min until the temperature of the water settling at the desired value (T= 15, 20, 25, 27, 29, 31, 33, 35). We put a single worm to the water and recorded a 5 min video in each temperature settings. **A.** We calculated the activity which is defined as a multiplication of change of the orientation ($\theta$) and position of a center of mass (CoM) (see inset of Fig.2B) as a function of the water temperature. To avoid transient response, we used second half of the recorded videos. The worms are most active at around 30 $^{o}$C. **B.** The major axis of the worms decreases with the temperature. The worms are more elongated in cold (around $\sim 20^{o}$C) water and started to shrink and curl up as the temperature increases (see Movie 2).

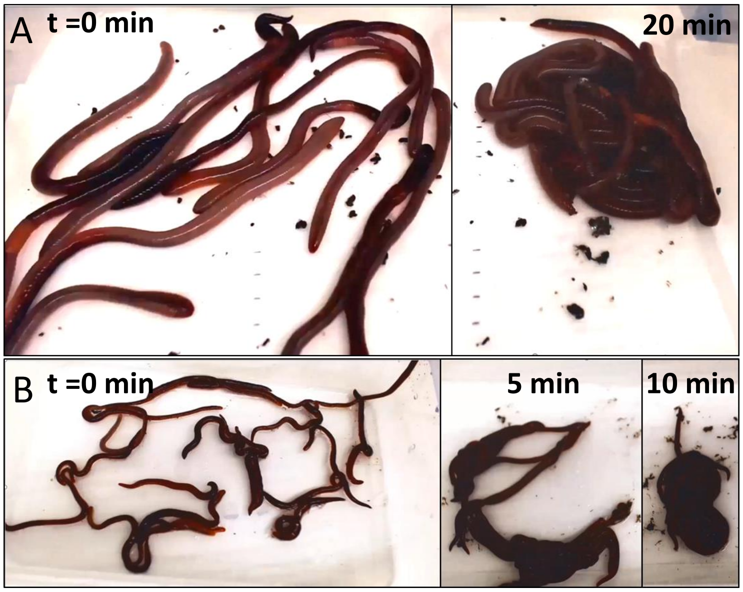


Figure 3: **Blob formation in other type of worms**– Blob formation is a stereotype behavior for other type of the worms. **A.** 13 earthworms (*Lumbricus terrestris*) with a length l= 19±2 cm were placed in a dry 20x30 cm box. They formed blob in 20 minutes. **B.** Same experiment was done with 23 red wigglers (*Eisenia fetida*) with a length l= ±2 cm. After 10 minutes they formed a blob (see Movie 2).

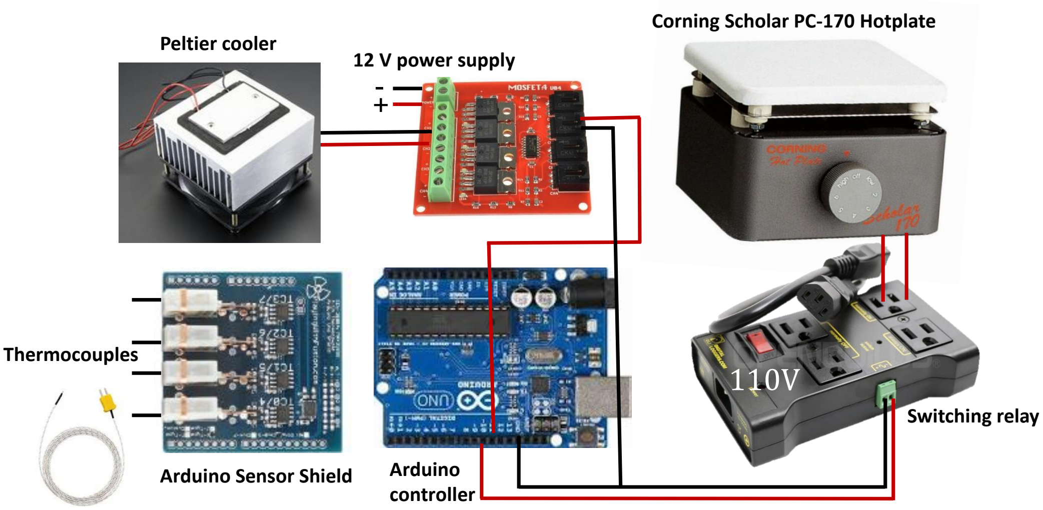


Figure 4: **The details of 3D thermal gradient setup**– To measure and control the temperature, two K-type thermocouples (Uxcell) were placed the two edges of the plate. Arduino sensor shield (playwithfusion.com, MAX31855 K-Type Thermocouple Sensor Breakout 4ch) used to collect data from the thermocouples. The peltier cooler (Adafruit) is swithed on/off with four channel MOSFET IRF540 and hot plate is switched on/off with switching relay(Openbuilds, IOT Switching Relay Power Strip), which are controlled by Arduino.

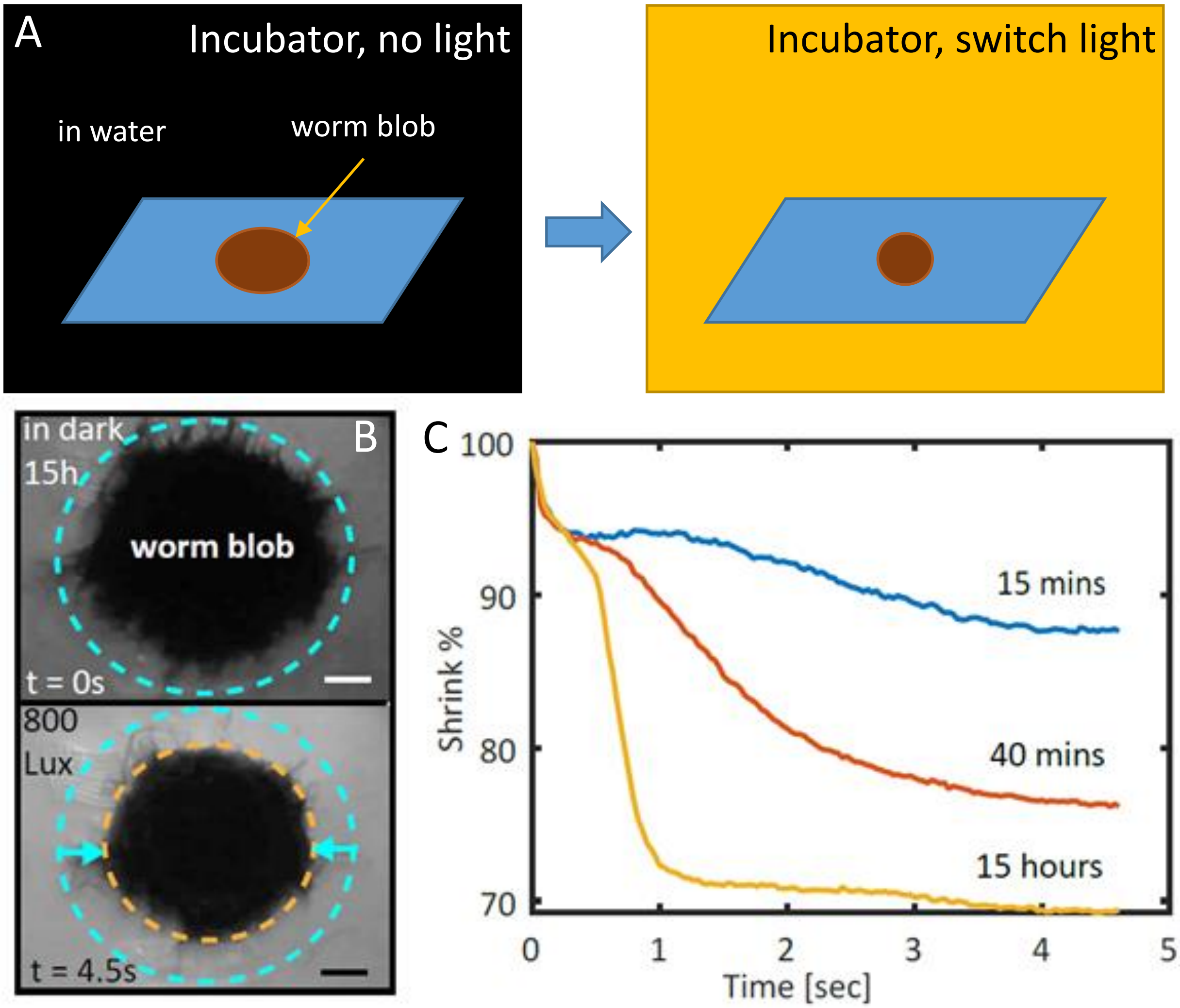


Figure 5: **Light response of the worm blob– A.** The worm blob was kept in dark in a temperature and humidity controlled incubator for a certain time and then we switched on the light. We measured the projected area of the blob for 5 sec. **B.** Phototaxis behaviour of the worm blob. Time snapshot from the experiment (top view) where worm blob were kept in the dark incubator for 15 hours and the light source (800 Lux ) was turned on. The blob shrank about 30% of its initial size. **C.** Phototaxis behaviour of the worm blob under different light stimulus history (duration of the time in the dark, see Movie 3) .

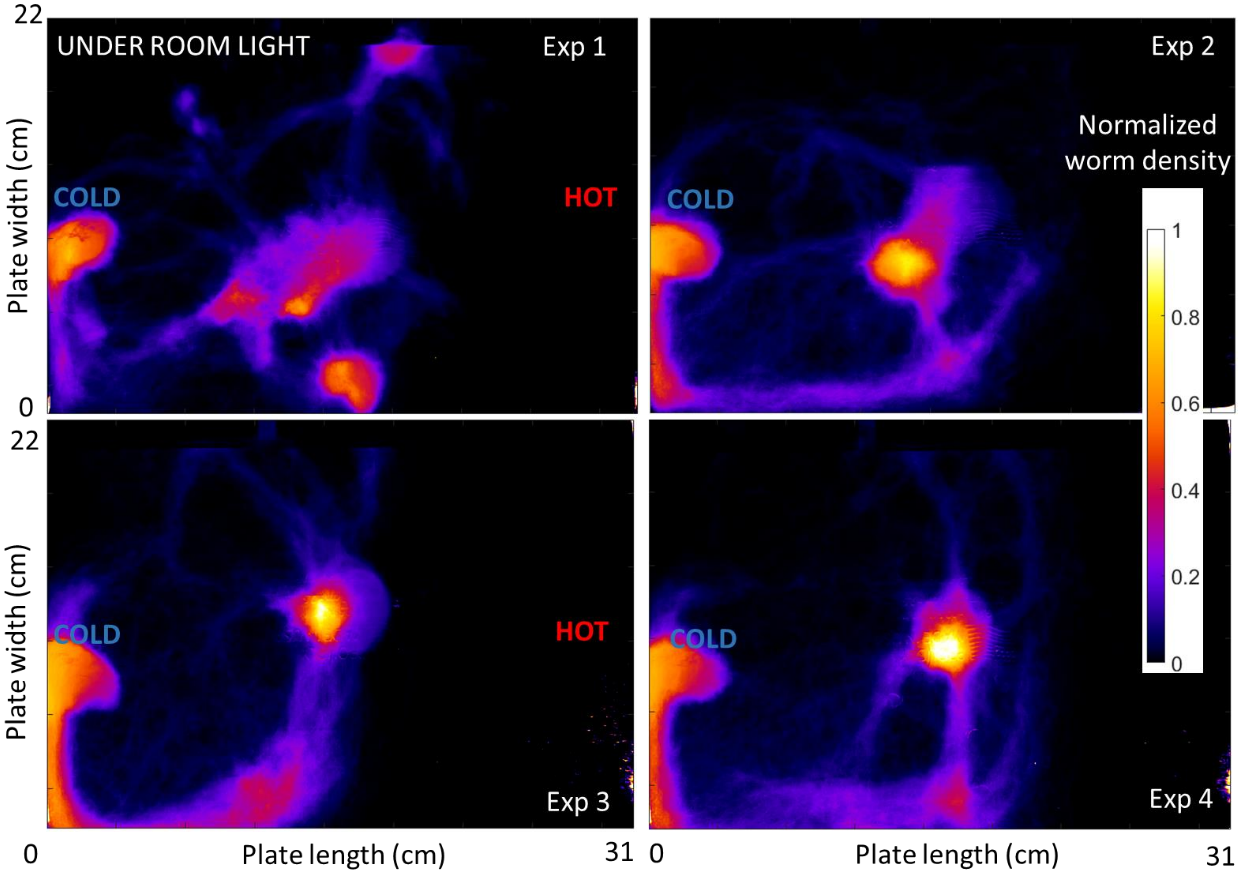


Figure 6: **Experimental space-time overlap heat maps of blob locomotion under ambient light**– The thermal gradient setup (left-cold, right-hot) used in this experiments is given in Fig.**??**A. At each time step, the density of the worms in the plate was calculated by counting black pixels and we plotted normalized pixel numbers at the end. Four panels show the results of four different experiments. The color bar represent normalized worm density (light color represents the most visited area). The heat map shows that the worms are moving individually.

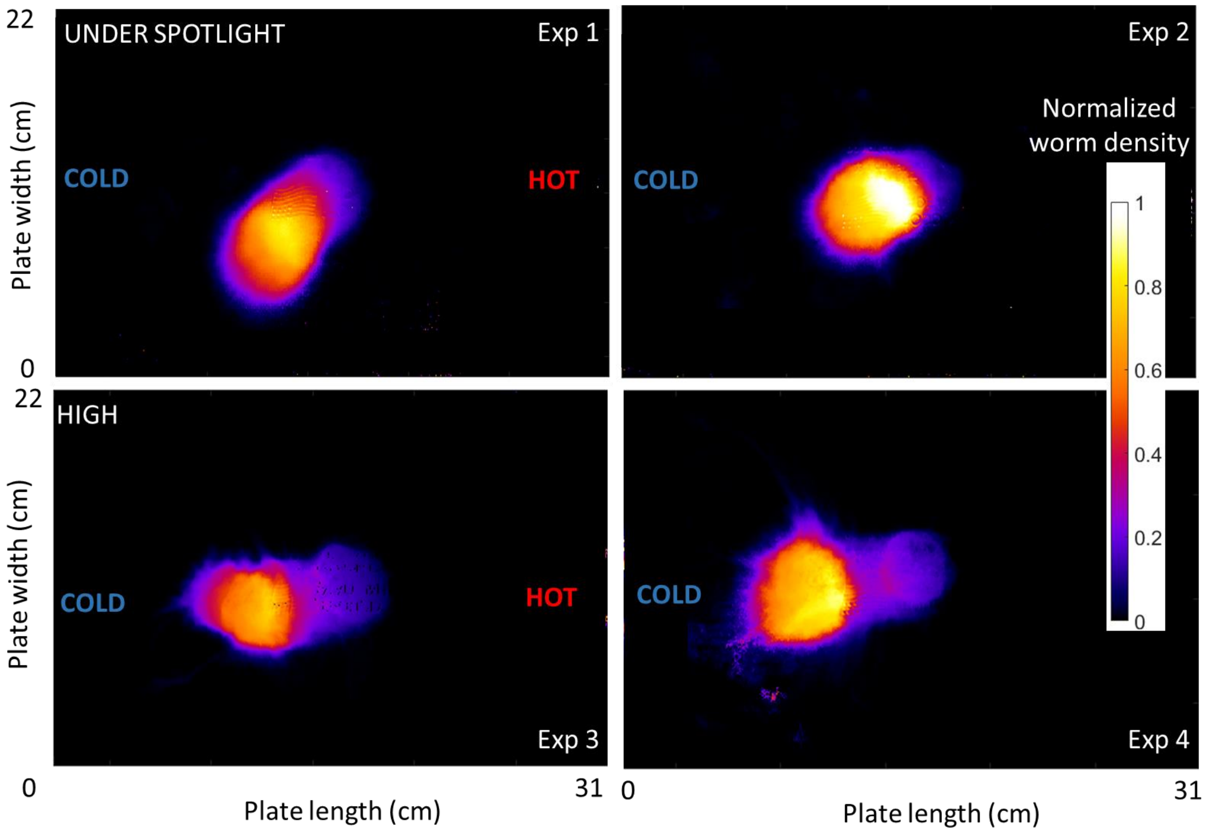


Figure 7: **Experimental space-time overlap heat maps of blob locomotion under spot light**– The thermal gradient setup (left-cold, right-hot) used in this experiments is given in Fig.**??**A. At each time step, the density of the worms in the plate was calculated by counting black pixels and we plotted normalized pixel numbers at the end. Four panels show the results of different experiments under 1500 Lux (top row) and 5500 Lux (bottom row) light intensity. The color bar represent normalized worm density (light color represents the most visited area). The heat map shows that the blob is moving as a group.

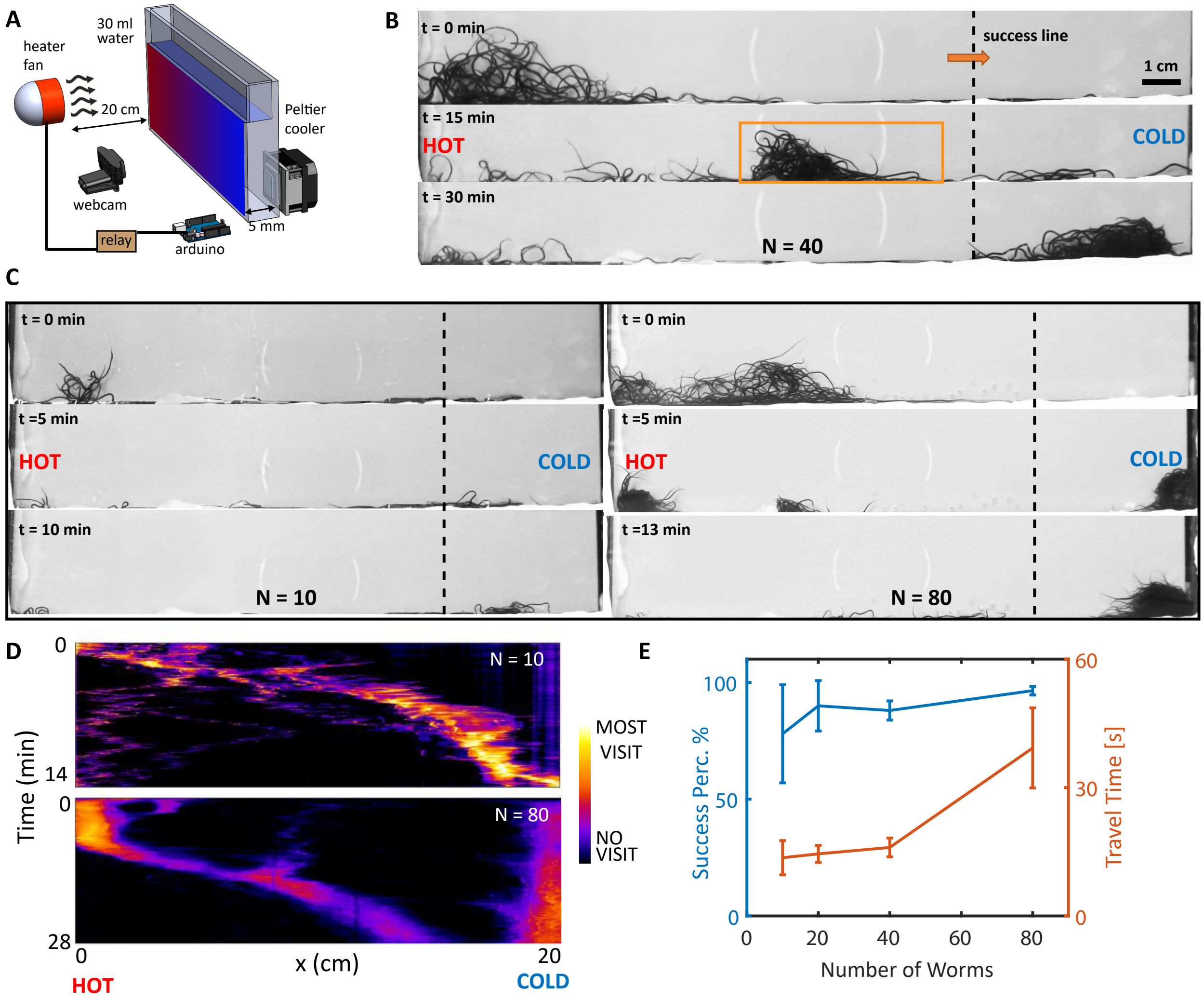


Figure 8: **Quasi 2D thermal response of the worm blob.** **A.** Experimental setup for quasi-2D thermal gradient experiments. The heater fan placed 20 cm away from the other side was activated for 10 seconds in every minute with an Arduino controller to create dynamic temperature gradient through the box. **B.** Time snapshot from the quasi-2D experiment (N = 40 worms). Right side of the dashed line shows the safe area used for calculation of the success percentage. **C.** Time snapshot from the quasi-2D experiment (N = 10 (left) and N = 80 (right) worms). **D.** Space-time overlap heat maps of blob locomotion (N = 10 (top) and N = 80 (bottom)). At each time step, the density of the worms in the plate was calculated by counting black pixels and we plotted normalized pixel numbers at the end. The color bar represent normalized worm density (light color represents the most visited area). **E.** Travel time (blue) and success percentage (red-defined as the percentage of the worms that craw to cold side) vs. number of worms (10, 20, 40 and 80), averaged over 5 experiments per each size.

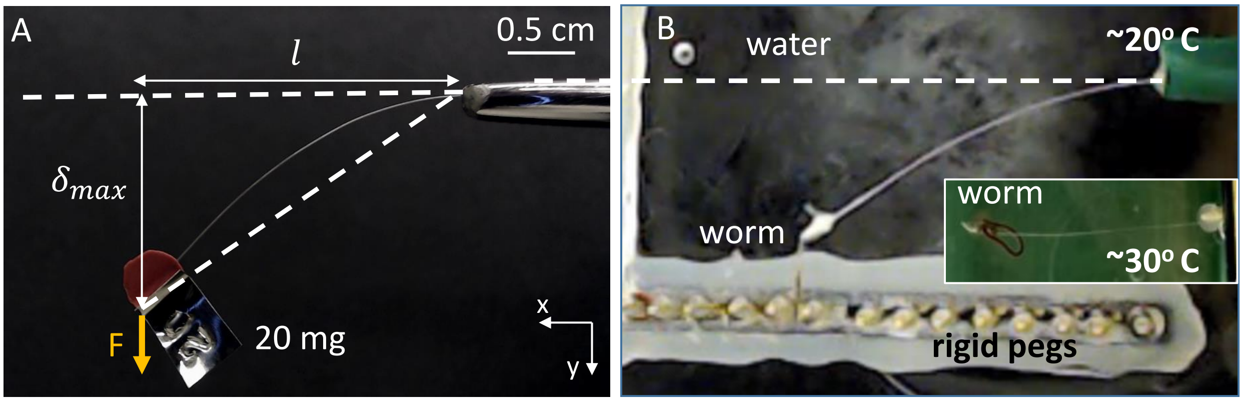


Figure 9: **Single worm pulling force measurement in different temperature–A.** We calculated the flexural rigidity (EI) of a stainless steel (l=3 cm, d = 0.1143 mm, A-M Systems, Inc., WA ) beam by applying known perpendicular external loads (F) to the tip of the beam (see Movie 4). The external force generated by the worm then can be calculated by the deflection formula $F = \frac{\delta_{max} 3EI}{l^3}$, where $\delta max$ is the maximum deflection of the tip of the beam **B.** Snapshot from one of the experiment where the worm apply maximum pulling force in cold water ($\sim 20^o$). Inset shows the curled shape of the worm in hot water ($\sim 30^o$).
